# 2.5D Actuating Substrates Enable Decoupling the Mechanical and Biochemical Effects of Muscle Exercise on Motor Neurons

**DOI:** 10.1101/2024.03.02.583091

**Authors:** Angel Bu, Ferdows Afghah, Nicolas Castro, Maheera Bawa, Sonika Kohli, Karina Shah, Brandon Rios, Vincent Butty, Ritu Raman

## Abstract

Emerging *in vivo* evidence suggests that exercise impacts peripheral nerves, but the difficulty of isolating and studying the muscle-specific impact on motor neurons *in vivo*, as well as the inability to decouple the biochemical and mechanical impacts of exercise in this setting, motivate investigating this phenomenon *in vitro*. In this study, we show that tuning the mechanical properties of fibrin hydrogels can generate stable 2.5D motor neuron and contractile skeletal muscle cultures that enable long-term efficient secretome harvesting from exercised tissues. Motor neurons stimulated with muscle-secreted cytokines significantly upregulate neurite outgrowth and migration, with an effect size dependent on exercise intensity. Actuating magnetic microparticles embedded within 2.5D substrates enabled us to dynamically stretch motor neurons and non-invasively mimic the mechanical effects of exercise, revealing that dynamic stretch has an equally significant impact on axonogenesis. RNA sequencing revealed different transcriptomic signatures between groups, with biochemical stimulation having a significantly greater impact on cell signaling related to axon growth and development, neuron projection guidance, and neuron-muscle synapse maturation. Our study thus leverages 2.5D actuating substrates to robustly validate a hypothesized role for muscle exercise in regulating motor neuron growth and maturation through both mechanical and biochemical signaling.

## 1. Introduction

Exercise has systemic beneficial effects on the body.^[3,4]^ Significant evidence suggests that exercise can impact the healthy morphology and function of a range of tissues including muscle, bone, fat, vasculature, immune cells, and the central and peripheral nervous systems, but the mechanisms by which physical activity regulates inter-tissue communication are still poorly understood.^[5]^ A growing body of literature, largely in animal models, has provided compelling evidence that exercise upregulates secretion of muscle-originating cytokines, often termed “myokines”, which are released into the circulatory system and can modulate cell signaling throughout the body.^[6–9]^ However, as many other cell types can secrete these cytokines, often in an exercise-mediated manner, it is difficult to isolate the muscle-specific origin of circulating biochemicals in the complex multicellular *in vivo* environment, or to understand their specific biological impact.^[10,11]^ Furthermore, while most previous studies have largely focused on the biochemical impacts of exercise, a few have also recognized that the large localized mechanical forces generated during muscle contraction have a mechanobiological impact on surrounding tissues such as tendon and bone.^[12,13]^ However, current experimental techniques do not readily enable isolating and studying the mechanical impact of muscle movement on specific cell types *in vivo*. Taken together, the literature on exercise thus highlights a critical need to advance fundamental understanding of the underlying biochemical and mechanical mechanisms by which muscle contraction influences intercellular signaling with other cell types.

While exercise-mediated intercellular crosstalk is relevant across many cell types, several recent studies have highlighted a particular need to study communication between skeletal muscle and motor neurons, since these two cell types work together to coordinate all voluntary movement.^[14–16]^ For example, we recently implanted a tissue engineered muscle graft into a mouse model of traumatic limb injury, and studied how targeted stimulation of muscle contraction within the graft, or localized exercise, impacted functional recovery. We observed that stimulated muscle grafts completely restored mobility to injured mice within 2 weeks of injury, and a phosphoproteomic analysis of the grafts revealed that exercise upregulated several cell-signaling pathways related to axonogenesis, neurite guidance, and neuromuscular junction formation.^[17]^ Our observation corroborated similar studies by others, highlighting that *in vivo* exercise training can enhance muscle innervation, and is correlated with upregulated circulating neurotrophins, such as ciliary neurotrophic factor, glial cell line-derived neurotrophic factor, and brain-derived neurotrophic factor.^[6,7,18,19]^ While these results indicate a potential role for muscle contraction in governing motor neuron growth, the difficulty of deconvolving the muscle-specific role of exercise in an *in vivo* context motivates investigating these questions in more tightly controlled *in vitro* environments. Moreover, developing and leveraging representative *in vitro* model systems for investigating exercise-mediated intercellular crosstalk would also enable, for the first time, decoupling the biochemical and mechanical impacts of muscle contraction on motor neuron growth and development.

Several interesting studies have mapped the secretome of engineered skeletal muscle tissues *in vitro*, and even shown that cytokines secreted from 2D muscle monolayers can be efficiently harvested to study signaling with other tissues such as bone.^[20–25]^ While some of these studies involve electrical pulse stimulation or mechanical stretch stimulation of the muscle cells,^[21,25]^ they do not demonstrate functional contraction of the muscle and are thus not fully representative of exercise, which requires sustained cyclic muscle twitch. It is likely that harvesting the secretome of exercising muscle tissues has proven difficult because 2D muscle monolayers tend to delaminate from hard tissue culture substrates after a few days in culture due to the large passive tension forces they generate during development and even larger active tension forces they produce during contraction.^[26–28]^ By contrast, 3D muscle tissues grown in soft extracellular matrix-mimicking gels are more representative of the *in vivo* mechanical environment and can be sustained for weeks or months in culture and exercised through repeated stimulated contraction.^[29–32]^ However, as 3D tissues typically require being cultured in large media volumes and have a much lower ratio of exposed cell surface area to tissue volume than 2D tissues, these systems are not ideal for harvesting secreted cytokines at high concentration. Moreover, as 3D muscles typically require being cultured in extracellular matrices containing Matrigel (a naturally-derived matrix with poorly-defined biochemical composition and significant batch-to-batch variability)^[33]^, it is difficult to isolate the impacts of muscle-secreted cytokines from growth factors being released by the Matrigel itself.

There is thus a significant need for robust *in vitro* platforms that enable long-term culture of exercised muscle in a Matrigel-free monolayer format that permits efficiently collecting secreted cytokines and investigating the biochemical impacts of muscle exercise on other cells, such as motor neurons. In addition, an *in vitro* platform that enables separately studying the mechanical impacts of exercise on motor neurons, which directly feel the forces generated during muscle contraction *in vivo*, would advance fundamental understanding of the multi-modal mechanisms by which exercise mediates muscle-nerve crosstalk.

In this study, we show that carefully tuning the stiffness of an extracellular matrix-mimicking hydrogel, fibrin, can be leveraged to generate stable contractile 2.5D muscle cultures that enable long-term efficient secretome harvesting from exercised tissues. We observed that our 2.5D platforms also served as effective substrates for monitoring motor neuron differentiation and growth, and that neurons biochemically stimulated with muscle-secreted cytokines significantly upregulated the total number, length, and rate of projecting neurites, as well as their total migration area, with an effect size dependent on the intensity of muscle exercise. Furthermore, we modified our recently established methodology for magnetic matrix actuation (MagMA)^[1,2]^ to show that magnetic microparticles embedded within 2.5D fibrin hydrogels could be actuated via a permanent magnet to impose mechanical forces on motor neurons in a non-invasive manner. These studies revealed that mechanically stimulating motor neurons by mimicking the stretch generated during muscle exercise upregulated neurite number, length, and rate of growth, as well as migration area by a similar amount as biochemical stimulation, even in the absence of any biochemical or biophysical contact cues from muscle. Despite morphological similarities between biochemically and mechanically stimulated neurons, RNA sequencing analysis revealed different transcriptomic signatures in response to both modes of signaling, with biochemical stimulation have a significantly greater impact on cell signaling related to axon growth and development, neuron projection guidance, and neuron-neuron and neuron-muscle synapse maturation.

Our actuating 2.5D matrix platform enabled robust *in vitro* validation of a previously hypothesized role for exercise in mediating muscle-nerve crosstalk, which has historically been challenging to mechanistically prove *in vivo*. Moreover, our methodology enables decoupling the different mechanisms by which exercise mediates nerve growth, showcasing the importance of isolating and separately studying the biochemical and mechanical impacts of muscle contraction on motor neuron growth and development. These results advance fundamental knowledge of intercellular signaling between skeletal muscles and motor neurons during exercise, and set the stage for future studies that leverage exercise as a tool to modulate neuromuscular intercellular signaling in physiological and pathological states. The 2.5D actuating substrates we have developed can, moreover, be used to decouple biochemical and mechanical crosstalk between muscle and other cell types in future, enabling deeper understanding of exercise-mediated intercellular signaling in the body.

## 2. Results

### 2.1. Designing 2.5D Substrate for Stable Long-Term Secretome Harvesting from Exercised Muscle

Strategies for extending the lifetime of skeletal muscle monolayers by preventing tension-induced delamination have largely focused on: 1) biochemical methods: coating tissue culture surfaces with proteins that encourage cell adhesion; and 2) mechanical methods: engineering substrates with microscale topography that enable stable cell-substrate tethering. The first of these methods involves depositing a thin layer of proteins such as gelatin, fibronectin, or collagen on top of plastic or glass tissue culture plates.^[26]^ These coatings mimic the biochemical environment of the native extracellular matrix and enable differentiating multinucleated muscle fibers from proliferating myoblasts in 2D cultures. However, long-term cultures of mature contractile muscle in this format are still precluded by issues related to cells peeling off their underlying substrate, likely due to the mechanical mismatch between the cells and substrate. We validated previous studies by testing the stability of muscle grown on gelatin coated-substrates at different concentrations (0.5% and 2% w/v) and for different times (5 min and 30 min). We observed that in all conditions, while differentiated muscle fibers were formed within 5 days of culture, significant regions of 2D tissue delaminated from the surface by Day 7, even in the absence of spontaneous or stimulated muscle contraction (Figure S1). Other groups have investigated alternative “2.5D” approaches to protein surface coatings by modulating the microscale topography of substrates to encourage stable adhesion of muscle monolayers. For example, a recent study demonstrated that muscles differentiated on polymeric nanofibrous substrates coated with Matrigel remain stably adhered up to 20 days and that the cell-substrate interface withstands muscle contraction.^[34]^ Likewise, other researchers have shown stable long-term muscle culture on a flexible silicon micro-cantilever coated with elastin, collagen, heparan sulfate proteoglycan, and hyaluronic acid.^[35]^ While these techniques are promising and better mimic the biochemical signals, mechanical environment, and physical topography of native extracellular matrices, they require complicated multi-step microfabrication and surface functionalization techniques to promote and maintain stable muscle adhesion.

Inspired by our established protocols for engineering mature stable 3D muscles within fibrin-based extracellular matrix hydrogels,^[36]^ we decided to test whether a thick fibrin gel with tunable stiffness could promote stable adhesion of a contractile muscle monolayer to a standard tissue culture plate using a 1-step fabrication process. Since these 2.5D substrates would provide cells with: 1) a biochemical environment that promoted cell adhesion and 2) a compliant mechanical environment that could withstand repeated muscle contraction, we anticipated that this method would leverage the positive aspects of both protein coatings and microfabricated substrates. Moreover, since fibrin gels can be easily and quickly formed by mixing liquid solutions of fibrinogen monomers and thrombin crosslinkers, and readily molded within any tissue culture plate, generating tunable substrates using this approach would not require complicated manufacturing processes.

Optimizing a fibrin gel substrate specifically for long-term skeletal muscle culture required first characterizing how the formulation of the hydrogel, specifically the concentration of the fibrinogen monomer and the thrombin crosslinker, impacted mechanical properties. While previous studies have investigated the impact of these variables on fibrin compressive modulus,^[37]^ rheological analysis of different fibrin formulations would enable a more representative characterization of the interactions between a contractile cell monolayer and its underlying substrate. We formulated fibrin gels of varying fibrinogen concentrations (4, 8, 12, 16 mg/mL) at a fixed thrombin concentration (0.4 U/mL), conducted a strain sweep to determine the linear regime (∼0.5% strain), and then performed a frequency sweep from 0 to 100 rad/s at this constant strain (**Figure 1**a-b). Comparing the elastic storage modulus, G’, at 10 rad/s (corresponding to ∼1 Hz, the frequency of muscle contraction used in our experiments) showed that G’ increased with increasing fibrinogen concentration (Figure 1c). By contrast, varying thrombin concentration (0.3, 0.4, 0.5 U/mL) at a fixed concentration of fibrinogen (8 mg/mL) had no significant impact on the storage modulus (Figure 1d), though we did observe that higher thrombin concentrations led to faster gelation timelines.

**Figure 1.**
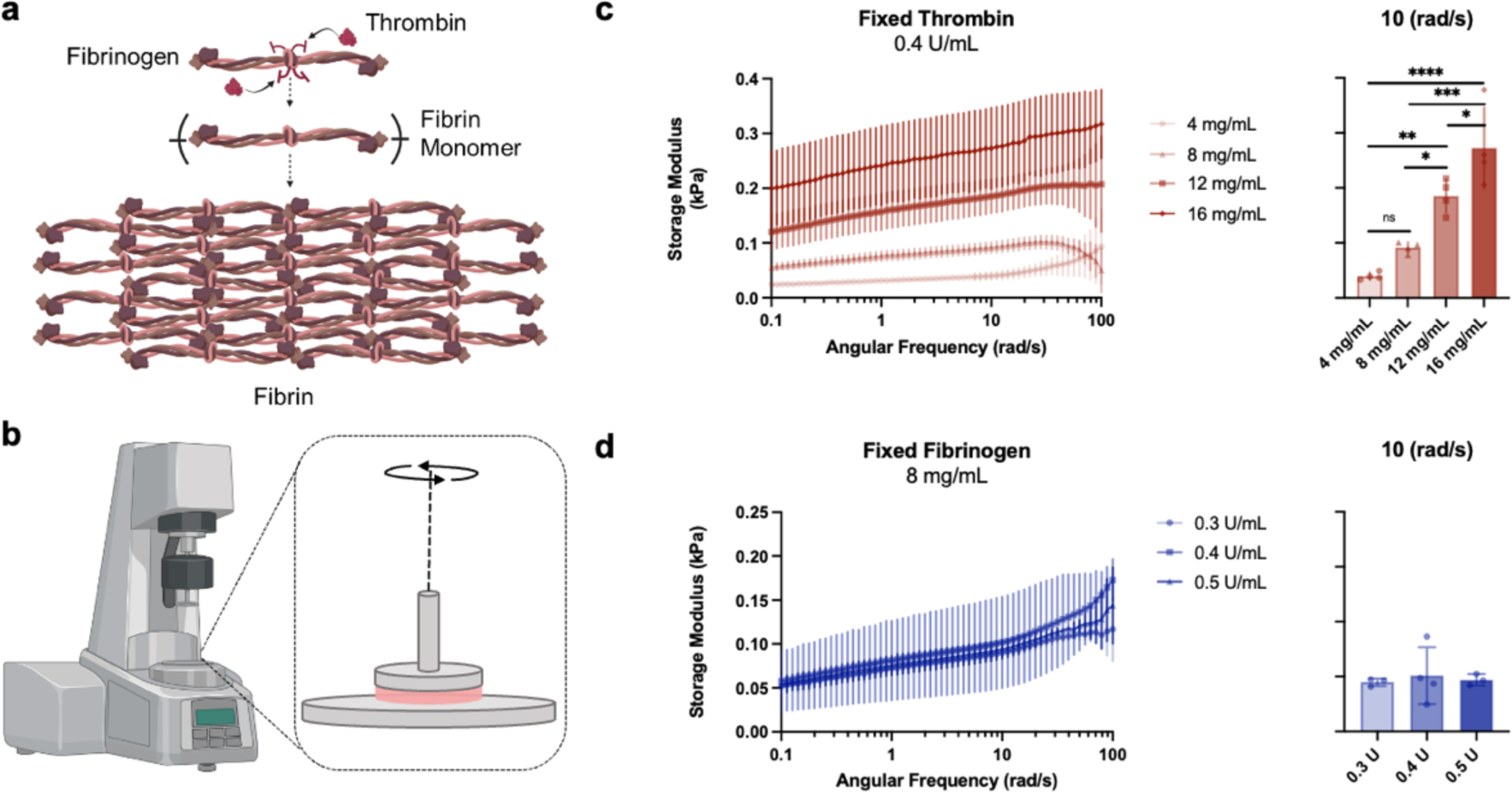
Optimizing fibrin hydrogel mechanical properties by varying fibrinogen and thrombin concentration. **a)** Schematic of fibrinogen and thrombin crosslinking to form a fibrin hydrogel. **b)** Schematic of our rheological testing setup. Made with BioRender. **c)** Varying the concentration of fibrinogen (4, 8, 12, 16 mg of fibrinogen per mL of solution) in a solution containing 0.4 U/mL of thrombin had a significant impact on the storage modulus of the gel. One-way ANOVA with Tukey’s multiple comparison tests, n=4 per group, p<0.05*, p<0.01**, p<0.001***, p<0.0001****. **d)** Varying the concentration of thrombin (0.3, 0.4 and 0.5 U of thrombin) in a solution of 8 mg/mL of fibrinogen did not significantly change the storage modulus of the fibrin hydrogel.

Due to the macromolecular structure of the fibrin hydrogels, visual observation may not provide an accurate assessment of their bulk structure. Hydrogels typically exhibit elastic behavior at low strain rates and low deformations, and this behavior (initially formulated and extended by Treloar and Flory as the rubber elasticity theory, RET), serves as a framework to elucidate the structural properties of hydrogels.^[38,39]^ Leveraging RET enables estimating a hydrogel’s average mesh size by using the elastic moduli obtained from frequency sweep measurements. This mesh size corresponds to the distance (Å) between two adjacent crosslinks providing an average porosity.^[40]^ The elastic motion of polymer chains between these crosslinks significantly affects the structural properties. It can thus be interpreted as an index of porosity and attributed to the pore size. This correlation can be calculated using Equation 1:

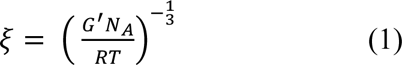

Where G’ is the storage modulus, N_A_ is the Avogadro number (6.022 × 10^23^), R is the gas constant (8.314 J/mol⋅K), and T is the temperature in K.

Deriving mesh size for varying fibrinogen concentrations showed that mesh size decreased with increasing fibrinogen concentration (**Table 1**). This observation can be attributed to the fact that higher fibrinogen concentrations lead to the formation of higher crosslinked gels characterized by denser structures and smaller pore sizes. Elevated fibrinogen concentrations likely promoted the establishment of larger networks with more branches, resulting in reduced distances between the crosslinking points and subsequently smaller mesh sizes.^[41]^

**Table 1.** Average mesh size for various fibrinogen concentrations derived from frequency sweep storage moduli using Equation 1. Consistent thrombin concentration of 0.4 U/ml applied across all experiments. Displayed are the mean values (n≥ 4) and corresponding standard deviations.

| Hydrogel concentration<br>(mg/mL) | $G'$<br>(Pa) | $\xi$<br>(nm) |
| --- | --- | --- |
| 4 | $38.33 \pm 5.65$ | $47.1 \pm 2.92$ |
| 8 | $101.44 \pm 7.9$ | $36.8 \pm 1.59$ |
| 12 | $170.29 \pm 22.0$ | $29.1 \pm 1.71$ |
| 16 | $256.81 \pm 20.5$ | $25.4 \pm 1.62$ |

For further testing with muscle monolayers, we selected an intermediate thrombin concentration with gelation timelines on the order of ∼2 minutes (0.4 U/mL) to enable easy molding into tissue culture plates, and the lowest fibrinogen concentration that demonstrated a stable elastic response even at high-frequency, 8 mg/mL. Modifying our established protocols for 3D skeletal muscle culture,^[36,42]^ we seeded optogenetic murine myoblasts (C2C12 engineered to express a 470 nm blue light-sensitive Channelrhodopsin [ChR2(H134R)]) on 1 mm thick fibrin gels cast in a standard 24-well plate format. Myoblasts were cultured in growth medium until they reached confluency and then transitioned to differentiation medium that encouraged cell fusion into mature multinucleated muscle fibers (**Figure 2**a).

**Figure 2.**
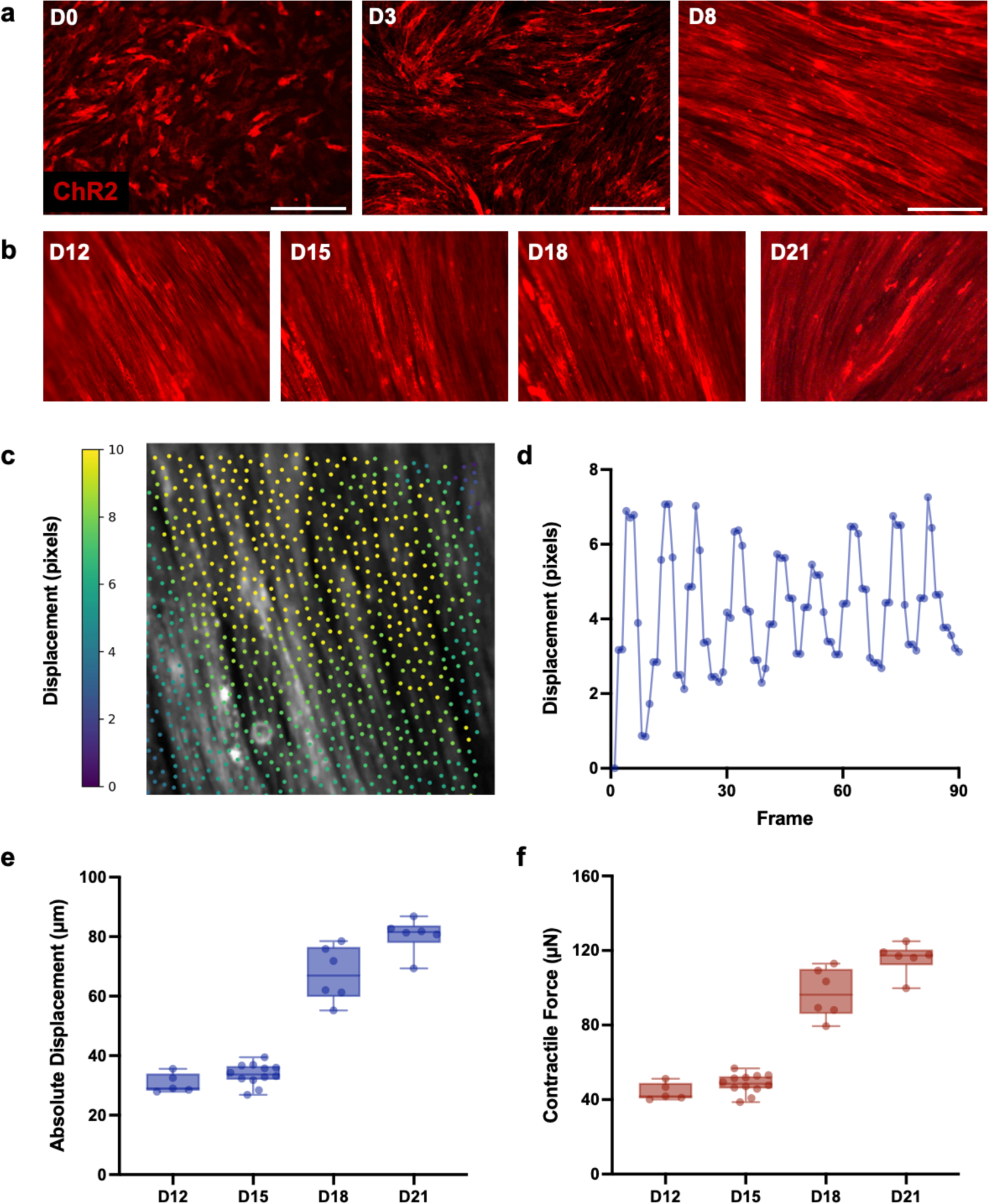
Longitudinal stability of contractile skeletal muscle in 2.5D fibrin culture format. **a)** Representative images of muscle tissue during early stages of muscle differentiation on days 0, 3, and 8. Scale bar = 300 µm. **b)** Representative images of muscle tissue after complete differentiation and observation of spontaneous twitch on days 12, 15, 18, and 21. **c)** Leveraging computational framework to spatially map muscle contraction on a single frame of a video (Video S1). **d)** Absolute displacement of representative 2.5D contractile muscle over multiple frames of a video. **e)** Peak absolute displacement of several 2.5D muscle tissues over multiple days in culture. **f)** Contractile force generated by 2.5D muscle tissue over multiple days in culture.

Differentiated muscle fibers started spontaneously twitching in culture within a week, indicating functional maturation of tissue. 2.5D tissues were exercised daily via light stimulation (1 Hz, 30 minutes) and remained stably adhered to fibrin gel substrates for the entire observation period we tested, 21 days (Figure 2b). We leveraged established open-source computational frameworks^[1,43,44]^ to convert videos of 2.5D muscle contraction into spatial maps of tissue deformation, and plotted muscle contractile displacements on Days 12, 15, 18, and 21 (Figure 2c-e, Video S1). As expected from previous 3D tissue studies, muscle spontaneous twitch force increased as tissues matured (Figure 2f). Importantly, the cell-matrix interface withstood repeated spontaneous contraction forces up to 125 µN without delamination, showing that 2.5D muscle tissues could generate forces comparable to those generated by 3D mm-scale muscle tissues (∼50-300 µN), despite being cultured on a

Matrigel-free matrix.^[29,42,45]^ These results indicated that our 2.5D platform was suitable for long-term stable cultures of highly contractile muscle monolayers, while still enabling efficient harvesting of the muscle secretome in response to exercise stimulation, as required for our proposed studies.

### 2.2. Impact of Biochemical Stimulation on Motor Neuron Growth

Fibrin gels have an established history of use as substrates not only for skeletal muscle, but also for motor neurons.^[46]^ Previous studies have shown that neurite outgrowth on fibrin gels is mediated by the stiffness of the substrate, with outgrowth being significantly upregulated by ∼2X on gels with low fibrinogen concentrations (< 10 mg/mL) as compared to gels with high fibrinogen concentrations (> 100 mg/mL).^[47]^ Moreover, as neurite outgrowth has been shown to be dependent on substrate stress relaxation rate,^[48]^ fibrin’s viscoelastic material properties make it a compelling substrate for motor neuron culture. Since the formulation of fibrin gels we optimized for maintaining contractile muscle monolayers utilized an 8 mg/mL fibrinogen concentration (i.e. less than 10 mg/mL), we decided to leverage the same substrate for monitoring the differentiation and growth of motor neurons. Preserving the same substrate for both tissues also enabled us to isolate the effects of our biochemical and mechanical stimulation experiments only to the specific method of stimulation we chose to test, rather than any effects related to differing substrate materials.

We followed established protocols to culture 3D neural spheroids from the HBG3 mouse embryonic stem cell line, and were able to visually confirm differentiation into a motor neuron phenotype since the cells expressed green fluorescent protein (GFP) linked to a motor neuron-specific promoter, Hb9.^[49,50]^ Following differentiation, motor neuron spheroids were seeded onto 2.5D fibrin gels and, over time, stably adhered to the substrate (**Figure 3**a-b).

**Figure 3.**
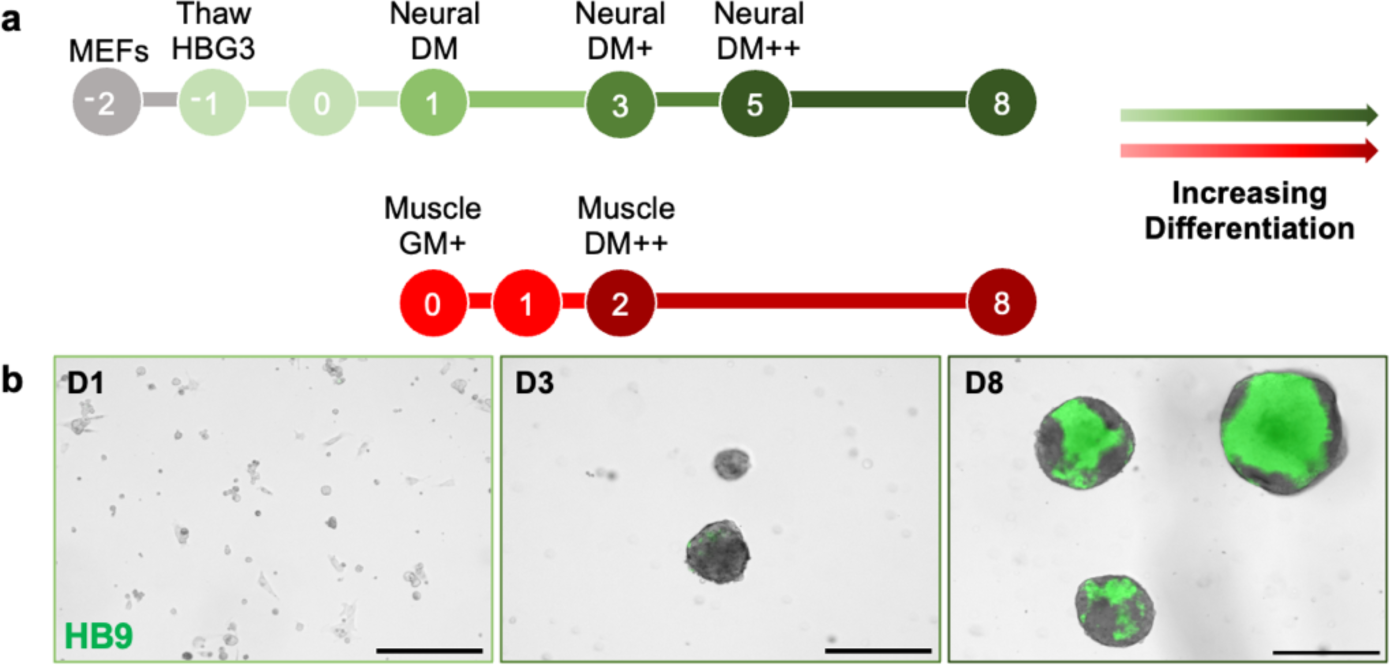
Cell culture timeline for HBG3 mESC motor neuron spheroids and C2C12 myoblasts. **a)** Timeline outlining the concurrent differentiation of motor neuron spheroids and skeletal muscle tissue. Motor neurons are grown on a substratum of mouse embryonic fibroblasts (MEFs) in neural differentiation media (Neural DM) supplemented with purmorphamine and retinoic acid (Neural DM+) and glial derived neurotrophic factor and ciliary neurotrophic factor (Neural DM++). Muscles are differentiated in muscle differentiation medium supplemented with aminocaproic acid and human insulin-like growth factor 1 (Muscle DM++). **b)** Representative images (fluorescent overlay on brightfield image) for HBG3 mESC on Day 1 showing no HB9 expression (green). On Day 3, the cells begin to form spheroids in the low adhesion petri dish and demonstrate slight HB9 expression. By Day 8, spheroids express HB9 throughout most of the spheroid, indicating successful differentiation to a motor neuron lineage. Scale bar = 300 µm.

We differentiated 2.5D muscle tissues in parallel with neural spheroids to study the biochemical effects of muscle exercise on motor neuron growth. To generate conditioned neuron medium containing a high concentration of muscle-secreted cytokines, we incubated muscle tissues in neuron medium and exercised the muscle monolayer. Specifically, we subjected the tissues to 470 nm light stimulation at 1 Hz for 30 minutes, following previously optimized training protocols from our 3D muscle studies.^[29]^ The exercise-conditioned neuron medium was then removed from the muscle culture and added to the motor neuron spheroid culture (**Figure 4**a). Observing response to biochemical stimulation with muscle-secreted cytokines over multiple days showed significant morphological differences between control and biochemically stimulated spheroids (Figure 4b). Quantifying standard metrics of neurite outgrowth over 5 days, namely the magnitude and rate of total neurite length, maximum neurite length, and neurite migration area, revealed significantly upregulated growth in exercised versus control groups across all metrics, with the exception of the rate of maximum neurite length (Figure 4c-e). Of particular interest, total neurite length, a metric of all the neurites projecting outward from the spheroid containing motor neuron soma, was increased by ∼3X in magnitude and ∼3.2X in rate after 5 days. Likewise, migration area of all projecting neurites was increased by ∼2.5X in magnitude and rate. These results showcased a significant impact of exercise-secreted cytokines on motor neuron growth.

**Figure 4.**
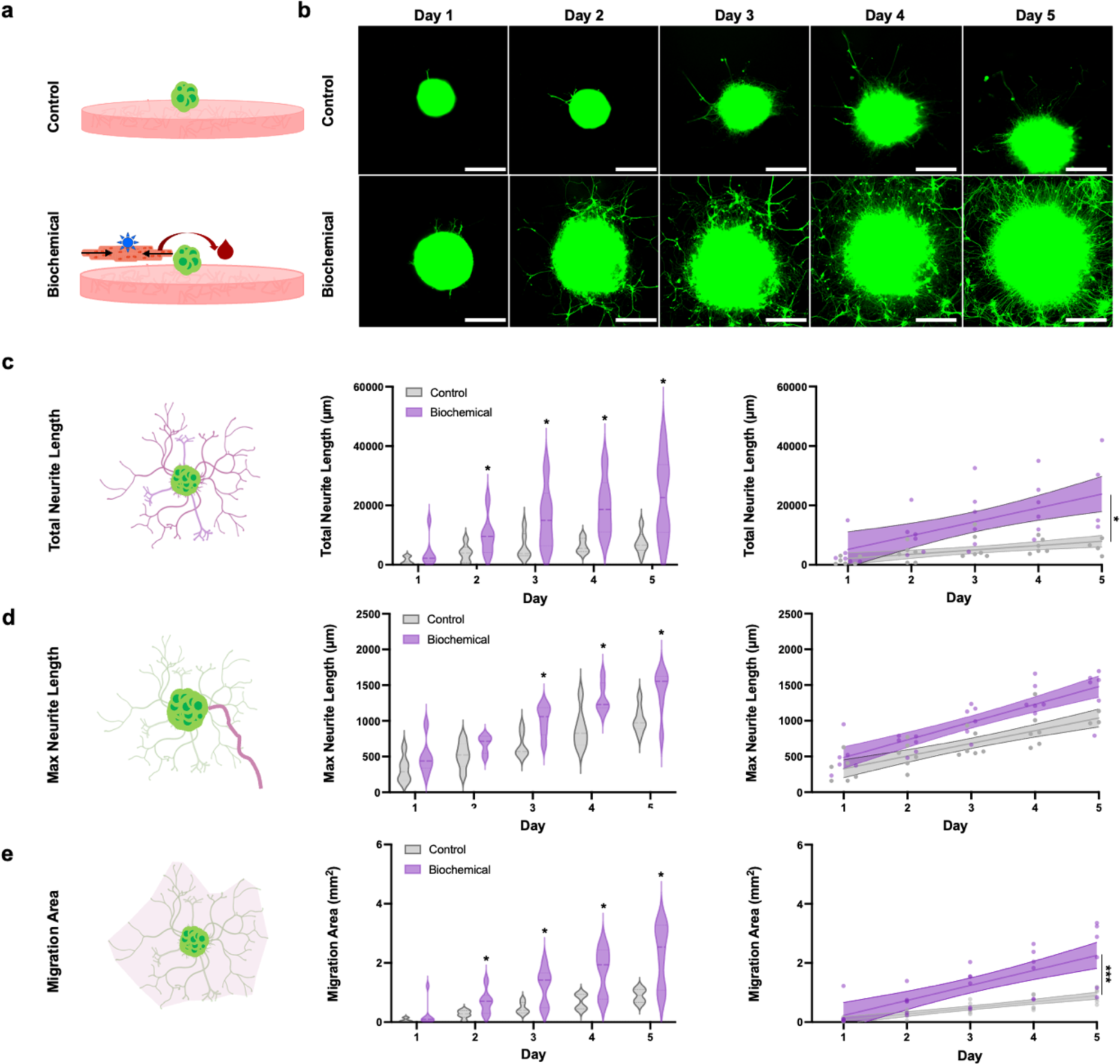
Impact of biochemical stimulation on motor neuron growth. **a)** Schematic of control and experimental group showing biochemical stimulation is conducted by supplementing motor neurons with conditioned media from exercised muscle. **b)** Representative images demonstrating motor neuron growth throughout the duration of the experiment. Scale bar = 300 µm. Biochemical stimulation was shown to significantly increase the **c)** total neurite length, **d)** maximum neurite length, and **e)** migration area in the later stages of the experiment. The rate of growth for total neurite length and migration area were also significantly impacted by biochemical stimulation. Unpaired t-test, n = 6 per group, p<0.05*, p<0.001***.

Since our 2.5D muscle tissues were spontaneously twitching through most of their differentiated lifetime, as is commonly observed by ourselves and others in mature *in vitro* muscle cultures, we decided to test the impact of conditioning neuron medium with secreted cytokines from muscles that were occasionally spontaneously twitching, but not actively exercised and paced by an external stimulus (Figure S2). These experiments revealed that even with this reduced level of muscle contraction, supplementing motor neurons with spontaneous twitch-conditioned medium significantly increased total neurite length, maximum neurite length, and neurite migration area. Notably, we observed that the relative morphological impact of biochemical stimulation was reduced in the spontaneous twitch (termed “low intensity exercise”) versus stimulated twitch (termed “high intensity exercise”) studies, with neurite length only being increased by ∼1.8X in magnitude and ∼1.5X in rate, and migration area being increased by ∼2X in magnitude and ∼1.7X in rate after 5 days. These data indicated that the effect size of biochemical stimulation on motor neuron growth is modulated by the amount of muscle contraction, lending further strength to the observation that exercise mediates biochemical signaling between muscle and motor neurons, likely in a dose-dependent manner.

### 2.3. Impact of Mechanical Stimulation on Motor Neuron Growth

In addition to being responsive to substrate stiffness, neurons are also responsive to dynamic mechanical forces in their environment.^[51,52]^ Studies have shown, for example, that imposing passive tension on motor neurons *in vitro* can significantly increase the rate of axon growth.^[53]^ *In vivo*, motor neurons extend from the spinal cord to distal skeletal muscle, and neurites navigate through dense muscle tissue to form physical connections with individual muscle fibers via neuromuscular junctions.^[15]^ Motor neurons and skeletal muscle are thus mechanically coupled in their native environment, highlighting the need to study whether and how the dynamic forces generated during muscle contraction regulate motor neuron growth.

We have recently developed a method for non-invasively mechanically stimulating cells termed magnetic matrix actuation (MagMA).^[1]^ Briefly, magnetic silicone microparticles embedded in an extracellular matrix hydrogel can be actuated by moving an external permanent magnet, thus yielding controllable deformation of the matrix (**Figure 5**a). Magnetically-enabled control of matrix actuation thus enables non-invasively imposing mechanical forces on the cells seeded on the hydrogel substrate. Leveraging our published data on how microparticle size and density and magnetic field strength modulate matrix deformation,^[1]^ we embedded a 2 *x* 3 array of magnetic microparticles (500 µm long by 100 µm wide by 100 µm tall) in our optimized formulation of 2.5D fibrin hydrogels, and leveraged a permanent neodymium magnet to generate deformation of the hydrogel mimicking the movement generated during muscle monolayer exercise (Figure 5b).

**Figure 5.**
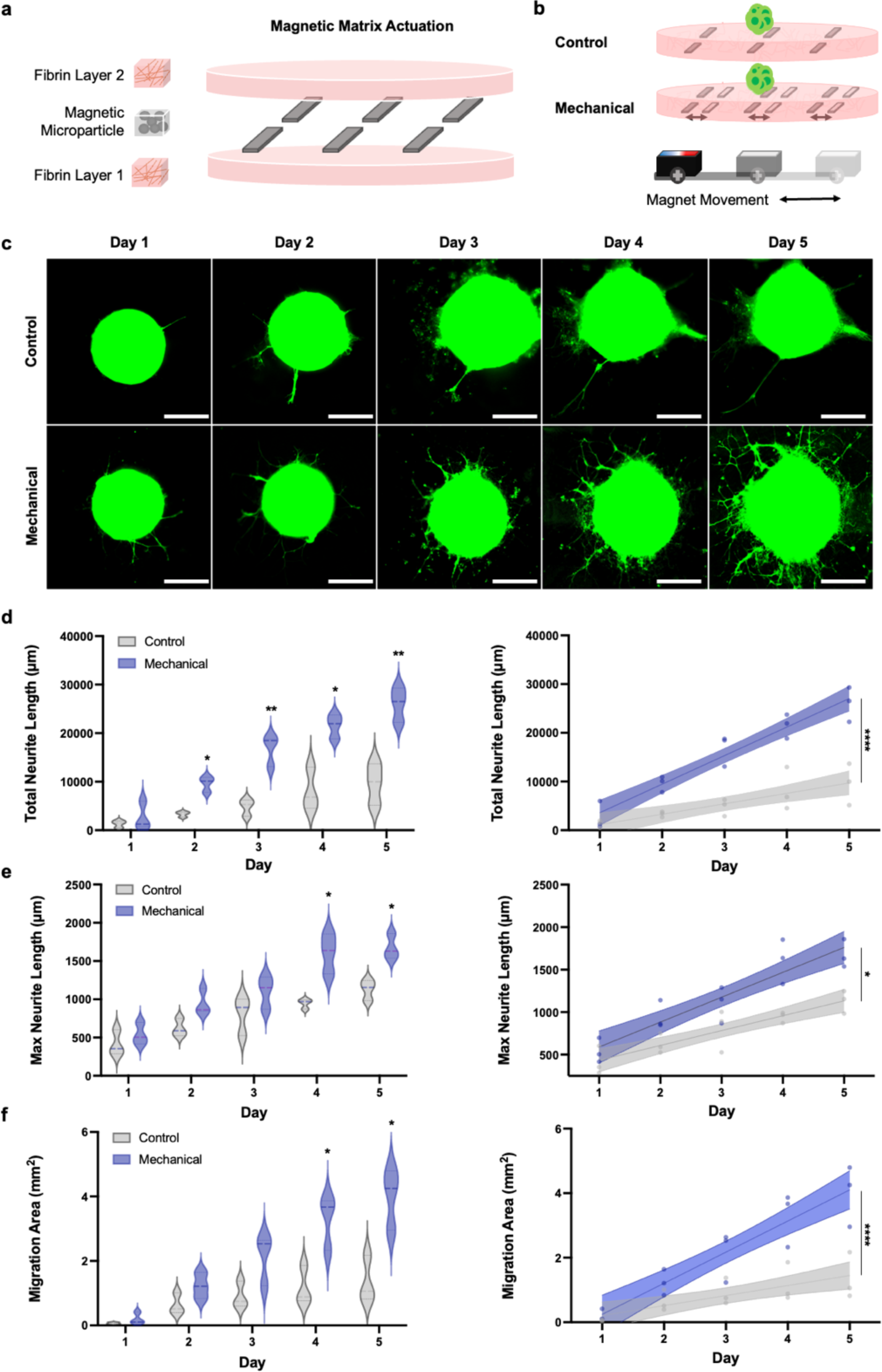
Impact of mechanical stimulation on motor neuron growth. **a)** Non-invasive mechanical stimulation of motor neuron spheroids was performed by leveraging our established methodology for Magnetic Matrix Actuation (MagMA). Magnetic microparticles embedded within fibrin hydrogels can be actuated using an external permanent magnet, thus deforming the substrate and mimicking the mechanical effects of muscle exercise. **b)** Both control and mechanical stimulation groups were seeded on MagMA fibrin hydrogels, but only spheroids in the mechanical stimulation group were actuated. **c)** Representative images for mechanically stimulated and control spheroids over 5 days. Scale bar = 300 µm. Mechanical stimulation was shown to significantly increase the magnitude and rate of growth of the **d)** total neurite length, **e)** maximum neurite length, and **f)** migration area. Unpaired t-test, n = 3 per group, p<0.05*, p<0.001***.

Motor neuron spheroids were generated as in previous experiments and seeded onto MagMA fibrin substrates and neurite outgrowth was monitored over a 5-day period in actuated (30 minutes daily) and non-actuated control hydrogels. Daily imaging revealed significant morphological changes in neural spheroids that were mechanically stimulated, as compared to controls (Figure 5c). Comparing the magnitude and rate of total neurite length, maximum neurite length, and migration area, as before, revealed significant increases across all metrics for mechanically stimulated neurons (Figure 5d-f). Specifically, after 5 days, total neurite length increased by ∼2.7X in magnitude and rate, and migration area increased by ∼3X in magnitude and ∼3.1X in rate, demonstrating a similar effect size in total neurite length and a greater effect size in migration area to motor neurons stimulated with conditioned media from high-intensity exercised muscle. These results indicated that mechanical stimulation of motor neurons during exercise may play an equally important role as biochemical stimulation in regulating neuron growth, though these effects have been largely neglected in prior animal studies due to the difficulty of isolating and studying mechanobiological phenomena in an *in vivo* context.

### 2.4. Transcriptomic Effect of Multi-Modal Exercise Stimulation on Motor Neurons

Our imaging studies outlined above enabled visualizing and quantifying motor neuron morphological changes in response to two modes of exercise-mimicking stimuli, and we were surprised to note similar effect sizes in response to biochemical and mechanical stimulation. To further investigate how different modes of stimulation impacted motor neuron maturation, we decided to perform RNA sequencing analysis of control and stimulated spheroids. We thus repeated the biochemical stimulation experiments outlined above and, after 5 days of stimulation, extracted RNA from the spheroids and studied the transcriptomic signatures of motor neurons in the high-intensity exercise, low-intensity exercise, and control conditions.

Principle component analysis of the three groups showed that both the exercised groups clustered together, and that the major variance between the biochemically stimulated motor neurons and the control non-stimulated motor neurons could be explained by 2 components (**Figure 6**a). Hierarchical clustering analysis of the three different groups again showed distinct clustering of control neuron spheroids as compared to biochemically stimulated neuron spheroids (Figure 6b). Interestingly, tissues in the high-intensity and low-intensity exercise groups also largely clustered within their own groups, with the exception of one high-intensity exercise stimulated-spheroid clustering with the low-intensity group. Differential gene expression analysis revealed 194 genes whose expression levels were significantly modulated in response to biochemical stimulation, as outlined in the volcano plots comparing gene expression between the three groups (Figure 6c-e). Pathway enrichment analysis showed significant modulation of pathways related to neuron projection development, axonal growth cones, and neuron-neuron and neuron-muscle synaptic signaling for both high-intensity and low-intensity groups, with effect size enhanced by high-intensity stimulation (Figure 5f-h).

**Figure 6.**
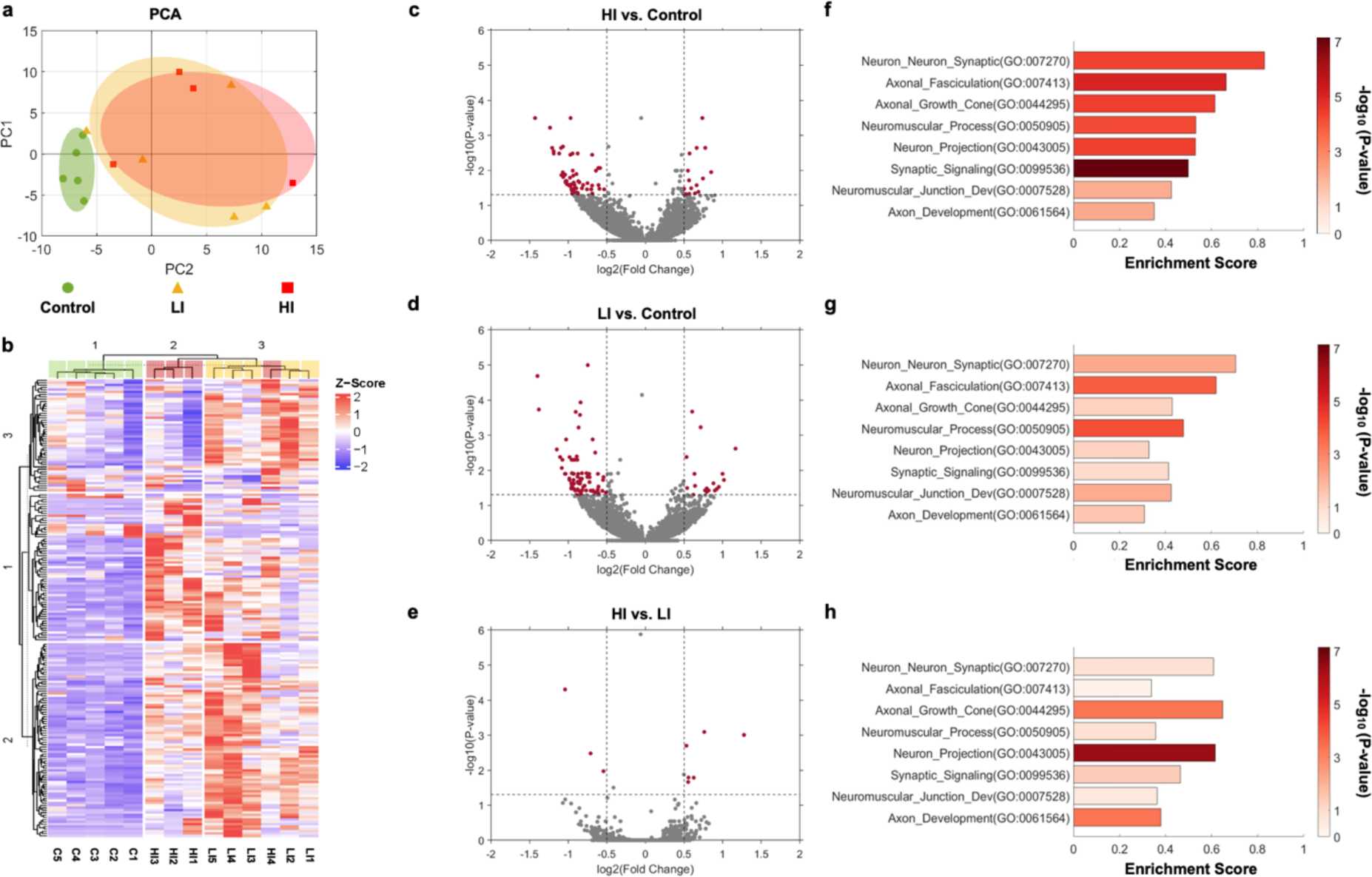
RNA-seq analysis of the impact of biochemical stimulation on motor neurons. **a)** Principal component analysis plot of our RNA-seq biochemical stimulation data. The control group had 5 biological replicates and the stimulation groups (low intensity exercise, LI and high intensity exercise, HI) had 4 biological replicates each. **b)** Heatmap showing all 194 differentially expressed genes with z-score normalization and hierarchical clustering. Volcano plots shows the differential gene expression between two groups: **c)** HI/Control, **d)** LI/Control, and **e)** HI/LI. Gene Set Enrichment Analysis (GSEA) results comparing the 3 groups: **f)** HI/Control, **g)** LI/Control, and **h)** HI/LI showing normalized enrichment score and nominal p-value. The p-values used to determine differentially expressed genes in the volcano plot and heatmap were Benjamini-Hochberg adjusted.

We also conducted RNA sequencing analysis of control and mechanically stimulated neural spheroids seeded on MagMA fibrin substrates. Interestingly, despite morphological similarities between mechanically stimulated and biochemically stimulated motor neurons, we noted that the transcriptomic impact of mechanical stimulation on motor neurons was significantly less powerful than in response to biochemical stimulation. Indeed, the variance between the two groups could not be explained by only 2 components, and differential gene expression analysis unveiled only 2 hits whose expression levels were significantly increased in response to mechanical stimulation, both corresponding to markers of RNA degradation (**Figure 7**a-b). Importantly, pathway enrichment analysis still showed significant modulation of neuron projection and axon development pathways in mechanically stimulated spheroids (Figure 7c). While it is typical to conduct hierarchical clustering analysis using Benjamini-Hochberg adjusted p-values, we performed hierarchical clustering analysis of our data using nominal p-values to find the top 50 genes whose expression was most modulated by mechanical stimulation, even if the effect was not significant. These analyses revealed that the mechanically stimulated spheroids still clustered separately from control spheroids (Figure 7c).

**Figure 7.**
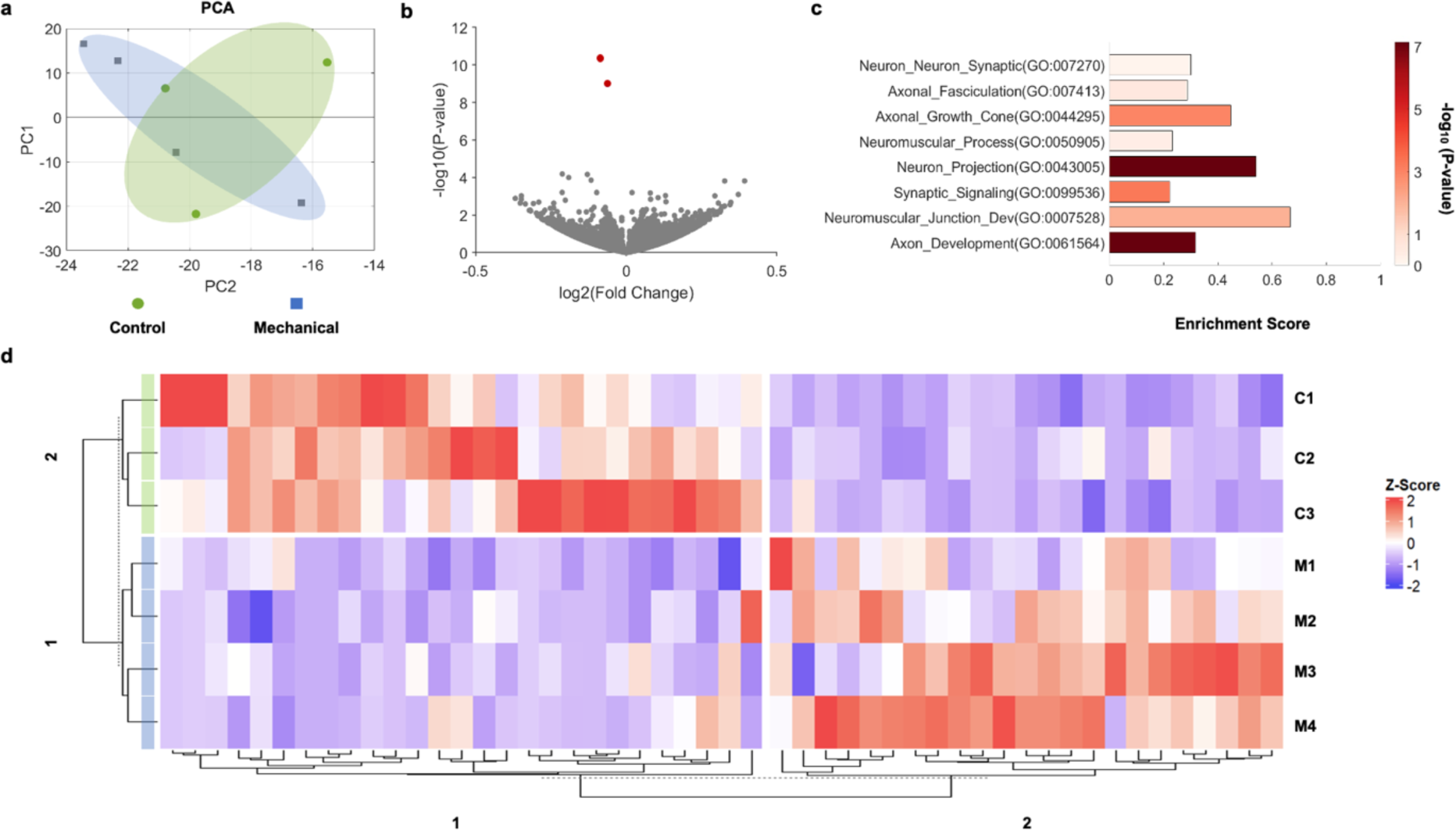
RNA-seq analysis of the impact of mechanical stimulation on motor neurons. **a)** Principal component analysis plot of our RNA-seq mechanical stimulation data. The control group had 3 biological replicates and the mechanical stimulation group had 4 biological replicates. **b)** Volcano plots showed that there were only two differentially expressed genes, and nominal p-values were plotted. **c)** GSEA results show minimal enrichment in the gene sets, but neuromuscular process and axon development were significantly different between the two groups. **d)** Heatmap of the genes with the 50 lowest nominal p-values, demonstrating that the two groups are distinctly discernible via hierarchical clustering.

Our results indicate that, in our study design, while both the mechanical and biochemical components of exercise have a significant and similar morphological impact on neurite outgrowth and migration area, mechanical stimulation has a less significant effect on the motor neuron transcriptome. Leveraging a 2.5D actuating extracellular matrix substrate thus enabled us to investigate and highlight the importance of decoupling the biochemical and mechanical effects of muscle exercise, and studying the separate mechanisms by which they modulate motor neuron growth and maturation.

## 3. Discussion

Despite significant *in vivo* data suggesting that muscle exercise regulates motor neuron growth and guidance, the difficulty of isolating and studying the muscle-specific role of exercise *in vivo*, as well as the inability to decouple the biochemical and mechanical impacts of exercise in this setting, motivate investigating this phenomenon further in relevant *in vitro* model systems. Given evidence from our own prior animal studies indicating that exercise upregulated cell signaling related to axonogenesis and neurite guidance,^[17]^ which aligned with emerging understanding in the literature of muscle as an endocrine organ,^[9]^ we aimed to design a representative *in vitro* model system for efficiently harvesting secreted myokines from exercised muscle tissues.

Measuring and optimizing the material properties of 2.5D fibrin gels enabled us to identify a formulation that provided biochemical and mechanical support to muscle monolayers and promoted robust long-term cell adhesion even in response to daily stimulated exercise. Leveraging optogenetic muscle cells, moreover, enabled us to exercise large batches of 2.5D tissues in a non-contact manner, thus avoiding potential downsides of gold-standard electrical stimulation approaches which can yield electrolysis of the culture media and damage tissues if used over extended time periods.^[42,54]^ We took advantage of our platform’s ability to efficiently generate large volumes of conditioned medium containing exercised muscle-secreted cytokines, and studied how supplementing motor neurons with conditioned medium impacted morphology. Corroborating previous *in vivo* hypotheses that had yet to be robustly proven *in vitro*, our data showed, for the first time, that myokines do in fact have a neurotrophic effect on cultured neurons, and significantly increased total neurite length, maximum neurite length, and migration area. Interestingly, we noted that the size of the neurotrophic effect was positively correlated with the intensity of exercise, aligning with our prior *in vivo* phosphoproteomic analyses of muscle showing that neurotrophic cell signaling pathways were altered in an exercise dose-dependent manner.^[17]^

We anticipated that mechanical stimulation of motor neurons, mimicking the movement that they may experience during muscle contraction, may also have an impact on their morphology. We thus modified our 2.5D gel substrate to include magnetic microparticles that could be actuated by an external magnet, thus imposing mechanical strains on motor neurons in a non-contact manner. Surprisingly, we noted that dynamic mechanical stretch (matched in magnitude and duration to our standard muscle exercise protocols) increased neurite length and migration area by a similar amount as biochemical stimulation. Despite the fact that the dynamic mechanical impact of exercise on motor neurons has rarely been investigated, to our knowledge, our findings indicate that this method of stimulation may have an equally significant impact on neuron growth and migration as biochemical stimulation.

Given the morphological similarities between motor neurons in biochemically and mechanically stimulated groups, we anticipated that RNA sequencing analyses of cells in these groups would reveal similarly significant differences between biochemically and mechanically stimulated and control tissues. This hypothesis was validated in neurons stimulated with conditioned media from both high-intensity and low-intensity exercised groups, with both modes of exercise being associated with significant modulation of pathways related to neuron growth, guidance, and maturation in a dose-dependent manner. Of note, neuron signaling related to neuron projection development and axonal growth cones was significantly more enriched in high-intensity versus low-intensity exercised groups. Surprisingly, differential gene expression analysis of the transcriptome of mechanically stimulated neurons and their controls showed essentially no difference between these groups. However, pathway enrichment analysis still showed that mechanical stimulation significantly modulated signaling related to neuron projection and axon development, though synaptic signaling pathways were notably less enriched than in biochemically stimulated groups.

We attribute these different transcriptomic signatures at least partially to our experimental setup. When our 3D motor neuron spheroids are cultured on fibrin gels, only the neurons directly in contact with the actuating substrate feel the impact of mechanical stimulation. By contrast, all the neurons within the spheroid are impacted during biochemical stimulation with conditioned medium from exercised muscle, as cytokines can easily diffuse throughout the whole tissue. Bulk RNA sequencing analysis of the whole spheroid may thus not be the best method for investigating the transcriptomic impact of different modes of stimulation. In future studies, we hope to leverage spatial RNA sequencing methods to more specifically study how gene expression varies in neurons that directly feel the impact of dynamic stretch. However, even with spatial RNA sequencing, it is possible that we will find that mechanical stimulation still may not have as broadly significant an effect as biochemical stimulation. Indeed, we have noted similar trends in research by other groups studying the impact of mechanical stimulation on cells of the nervous system. Specifically, a recent study on the effect of stretch stimuli on microglia noted that mechanical stimulation impacted the mode of cell migration, and upregulated the secretion of one pro-inflammatory cytokine, but did not otherwise broadly impact the secretome.^[55]^

It is important to note that we did not try a wide variety of mechanical or biochemical stimulation protocols (with different frequencies, magnitudes, and durations) in our experiment. It is possible that further optimization of both stimulation methods separately or in synergy can enhance the morphological and transcriptomic impact of exercise stimuli on motor neurons even further. Given previous studies showcasing the positive impacts of electrically stimulating neurons *in vitro*,^[56,57]^ moreover, it would also be interesting to study whether coupling mechanical and biochemical stimulation with electrical stimulation of motor neurons could further accelerate the rate of cell growth and maturation.

While we believe our study significantly enhances fundamental understanding of the different mechanisms by which exercise mediates muscle-nerve crosstalk, it is important to note that our current *in vitro* platform design does not enable studying electrical communication between these two cell types via neuromuscular junctions. Future studies that also consider synaptic signaling between motor neurons and skeletal muscle in a co-culture could add an even deeper understanding to the different mechanisms of intercellular signaling at play. However, given that it would be impossible to decouple biochemical and mechanical signaling between muscles and motor neurons in a co-culture system, isolating and studying these modes of signaling in mono-culture formats (as we did in this study) was a necessary first step to advancing fundamental knowledge of how motor neurons sense and respond to exercise.

We hope that our 2.5D actuating matrix platform, which can be manufactured in a simple process without special tools, can be leveraged by others in the field to study biochemical and mechanical crosstalk between muscles and a range of different cell types. It is important to note that the gel substrate formulation may need to be optimized for different muscle cell lines, as there are typically significant differences in the forces generated by muscle cells depending on their animal of origin (mouse, human, etc.) and source (primary, cell line, stem cell-derived).^[58,59]^ However, we anticipate that the rheological data we have gathered will provide other researchers with quantitative parameters to aid with designing stable contractile muscle monolayers from a variety of cell sources.

Another experimental design item to consider is the large batch-to-batch variation between stem cell-derived neurons. For example, we have noted morphological differences between motor neuron spheroids derived from different batches of differentiation, and accounted for this in our experimental design by comparing each experimental group to its own control, composed of spheroids from the same batch. We encourage other researchers to adopt similar practices when conducting studies of muscle-motor neuron crosstalk so that observed impacts of stimulation can be attributed to the method of stimulation, rather than inter-batch variability. Looking beyond motor neurons, given several recent interesting studies showcasing how mechanical stimulation impacts morphology and function of a range of tissues,^[60,61]^ we hope to see others leverage our platform to decouple the biochemical and mechanical impacts of exercise on tissues such as fat, bone, tendon, and blood vessels.

Our study is a first step towards mechanistically unraveling how repeated muscle contraction, or exercise, regulates motor neuron growth and maturation through both biochemical and mechanical modes of signaling. Future *in vitro* studies that optimize our 2.5D platform and leverage multi-modal stimulation could further advance understanding of how exercise cues can be used to control muscle innervation in healthy and pathological states. In the long-term, by leveraging established tools for *in vivo* biochemical stimulation and mechanotherapy,^[62–65]^ we hope our *in vitro* learnings will translate to effective therapeutic strategies that preserve and promote healthy muscle innervation and mobility.

## 4. Experimental Methods

### 4.1. 2.5D Actuating Substrate Fabrication

#### 4.1.1. Fibrin Formulation

The fibrin substrate was made using a concentration of 8 mg of fibrinogen from bovine plasma (Sigma Aldrich, F8630) per mL of GM+ to 0.4 U of thrombin (Sigma Aldrich, T4648) from bovine plasma. The solution was then pipetted into a glass bottom 24-well plate (Cellvis, P24-1.5H-N) at a volume of 250 µL to create a gel thickness of ∼1 mm in height. The fibrin was set in an incubator at 37 °C to gelate for 30 minutes and promptly hydrated using GM+.

#### 4.1.2. Magnetic Matrix Actuation

The Magnetic Matrix Actuation (MagMA) platform was designed to be a 2 by 3 array of 500 µm long by 100 µm wide by 100 µm tall magnetic microparticles spread evenly inside the fibrin gel. To manufacture this platform, a thin layer of fibrin gel (150 µL) was first pipetted into the glass bottom well plates. The gel was then allowed to set for 30 minutes at 37° C. Next, the microparticles were cut from a large sheet of cured 1:33 PDMS (Sylgard 184) loaded with 25% w/w of 4 µm diameter iron microparticles (US Research Nanomaterials). The microparticles were laid on top of the first layer of fibrin hydrogel and were subsequently covered with another layer of fibrin hydrogel solution, 250 µL. The fibrin hydrogel was set to gelate one final time and promptly hydrated using GM++.

### 4.2. Cell Culture

#### 4.2.1. C2C12 Myoblasts

Optogenetic C2C12 myoblasts^[36]^ were initially cultured in muscle growth medium (GM) to promote cell proliferation. GM consists of a base DMEM with 4.5 g/L of glucose, L-glutamine, and sodium pyruvate (Fisher Scientific, MT10013CV) with 10% fetal bovine serum (Sigma Aldrich, F2442), 1% L-glutamine (Fisher Scientific, MT25005CI), and 1% penicillin-streptomycin (Fisher Scientific, MT30002CI). After expansion, the cells were seeded onto the prefabricated fibrin hydrogels at a density of 1.5e5 cells per well and suspended in GM medium plus 1 mg of aminocaproic acid per mL of growth medium (GM+). After 2 days on GM+ the cells are swapped onto our differentiation medium (DM++). DM++ is composed of the same base DMEM with 10% inactivated horse serum (Fisher Scientific, 26-050-070), 1% L-glutamine, 1% penicillin-streptomycin, 1mg/mL of aminocaproic acid, and 50 ng/mL of human insulin-like growth factor 1, IGF-1(Sigma-Aldrich, I1146). Differentiation medium was changed daily for the duration of the experiment.

#### 4.2.2. HBG3 Embryonic Stem Cells

We preplated a feeder layer of irradiated mouse embryonic fibroblasts in DMEM with 10% v/v of fetal bovine serum, 1% penicillin-streptomycin, and 1% v/v of L-glutamine the day before thawing our mESCs. Next, we seeded the HBG3 mESCs at a density of 5 E4 cells per cm^2^ onto the feeder layer on mESC proliferation medium. The proliferation medium consists of EmbryoMax ES DMEM (Sigma-Aldrich, SLM-220) with 15% v/v fetal bovine serum, 1% penicillin-streptomycin, 1% L-glutamine, 1% Embryo-Max nucleosides (Sigma-Aldrich, ES-008), 1% MEM non-essential amino acids (Thermofisher Scientific, 11140050), 0.1 mM β-mercaptoethanol (Thermofisher, 21985023), and 0.1% (v/v) ESGRO mLIF (Fisher Scientific, 26050070).

After 2 days, the mESC are fed neural differentiation media (Neural DM). After 1 hour, cells were trypsinized and resuspended in Neural DM on a standard tissue culture treated 10 cm dish. Neural DM is made up of a base of 50% v/v of Advanced DMEM/F12 (Thermofisher, 12634028) and 50% v/v of Neurobasal medium (Thermofisher, 21103049), and supplemented with 10% KnockOut serum replacement (Thermofisher, 10828010), 1% penicillin-streptomycin, 1% L-glutamine, and 0.1 mM β-mercaptoethanol. On the next day, we transferred the unattached cells onto a low-adhesion 10 cm petri dish. On day 3 of differentiation, we added 1 µM purmorphamine (Sigma-Aldrich, 540220) and 1 µM retinoic acid (Sigma-Aldrich, R2625) to the Neural DM (Neural DM+). After day 5 of differentiation, we added 10 ng/ml of glial derived neurotrophic factor (Neuromics, PR27022) and ciliary neurotrophic factor (Sigma-Aldrich, C3710) to the medium (Neural DM++). On day 8 of differentiation, the spheroids are ready to be transferred to the fibrin hydrogels.

### 4.3. Rheological Analysis

Dynamic mechanical properties of various formulations of hydrogels were characterized using a TA Discovery HR-2 Hybrid (TA Instruments) rheometer with a 20 mm parallel plate geometry at room temperature (25 °C). A gap of 1 to 2 mm was established, with a maximum axial force of 0.1 N. The experiments were repeated a minimum of four times and average data was presented.

The preparation procedure involved measuring specified quantities of fibrinogen reconstituted in GM+. The highest concentration was prepared first and then diluted into lower amounts. Thrombin was then added and mixed in. The hydrogels were rapidly poured into circular PDMS wells, each 25 mm in diameter and 4 mm in height, at equal volumes of 1 mL. Subsequently, the hydrogels were hydrated using culture media and incubated at 37 °C for 45 minutes to allow gelation. The final concentrations of hydrogels were 16, 12, 8, and 4 mg of fibrinogen per mL of cell culture media.

Samples were retrieved from the PDMS well, cut into smaller diameters using a 22 mm biopsy punch, and loaded onto the plate. A covering trap was employed to minimize water evaporation during the tests. Two sets of assessments were conducted: oscillatory amplitude sweep, and frequency sweep tests were performed on gels with different concentrations of fibrinogen and thrombin. The former aimed to ascertain the linear viscoelastic (LVE) region of the hydrogels. Storage (G’) modulus was recorded as a function of strain values at a constant angular frequency of 10 Hz, spanning a range from 10^−2^ to 10^2^% strain. The LVE range for each hydrogel concentration was chosen accordingly.

The viscoelastic properties of the hydrogels were then explored across a frequency range from 0.1 to 100 rad/s, maintaining strain values at 0.5 and 0.6% for fibrinogen concentrations of 4 and 8 mg/mL, and 12 and 16 mg/mL, respectively.

### 4.4. Biochemical Stimulation

Light stimulation was utilized to induce contraction in our mature contractile 2.5D muscle cultures. We 3D printed a housing compartment to fit directly over our 24 well plate with two circular openings on top to pulse two 5W blue LED bulbs. A function generator was used to pulse the light output at 1Hz, 20% duty cycle, for 30 minutes each session.

Before light stimulation begins, we aspirated the DM++ in the C2C12 monoculture and added our complete neural differentiation medium. After the muscle layer underwent light stimulation, we collected the media and centrifuged it to remove any floating cells in the media to pipette onto the neuron culture. DM++ was added to the wells after aspirating the stimulated media.

### 4.5. Mechanical Stimulation

To isolate the mechanical stimulation on neurons undergone during muscle contraction we utilized our MagMA platform. We placed our magnetic hydrogel above the linear actuator containing the permanent magnet. The daily exercise consisted of 30 minutes of 0.33 Hz of magnetic actuation. The well plates were alternated to keep external stresses similar across samples.

### 4.6. RNA Sequencing

RNA extraction was performed on motor neuron spheroids at the end of the experiments using a Qiagen RNeasy Micro kit following the manufacturer’s instructions.

The data discussed in this publication have been deposited in NCBI’s Gene Expression Omnibus^[66]^ and are accessible through GEO Series accession number GSE257540: https://www.ncbi.nlm.nih.gov/geo/query/acc.cgi?acc=GSE257540

#### 4.6.1. RNA-Seq Library Preparation

Sequencing libraries were generated using Takara’s SMARTR ZapR v2-based protocol; briefly, 4 ng of each RNA sample were used (corresponding to half of the reaction volume). Full volume was used for the 8 samples with the lowest concentration. RNAs were fragmented for 4 minutes, followed by first-strand cDNA synthesis using the SMARTR Pico v2 protocol. TruSeq-Singular unique dual index (UDI) sequencing adapters were added by PCR (5 cycles). Library fragments corresponding to rRNA molecules were depleted by cleaving with the ZapR v2 and mammalian-specific R-probe v2 kit at full volume, and the second round of PCR was performed using the Singular qPCR primers for 15 cycles. Library sizes were quantified and verified by QPCR and on a Fragment Analyzer before loading for sequencing on a Singular G4 instrument in a 50-base paired-end configuration.

#### 4.6.2. RNA-Seq Analysis

Reads were aligned against mm10 (Feb., 2009) using bwa mem v. 0.7.12-r1039 with flags –t 16 –f, and mapping rates, fraction of multiply-mapping reads, number of unique 20-mers at the 5′ end of the reads, insert size distributions and fraction of ribosomal RNAs were calculated using bedtools v. 2.25.0.^[67]^ In addition, each resulting bam file was randomly down-sampled to a million reads, which were aligned against hg19, and read density across genomic features were estimated for RNA-Seq-specific quality control metrics. Fastq files were processed using the nf-core/rnaseq v 3.11.1 pipeline [https://zenodo.org/records/ 10171269] in the Nextflow 22.10.4 environment, using the GRCm39 reference genome and ENSEMBL GRCm39.110 murine annotation. Quality control metrics were inspected for consistency and samples with low read counts (< 500K reads), large fraction of ribosomal RNA (> 20%) or low exon/intron read coverage ratios (< 10) were excluded from further analysis (n = 5). Gene-level expression tables generated by STAR/Salmon processing were retrieved.^[68–70]^ Log-transformed transcript-per-million (TPM) values were used for data visualization. Differential gene expression was tested using DESeq2^[71]^ in the R v 4.2.0 statistical environment using both a normal prior and the ashr statistical procedure.^[72]^ Log2 fold-changes, raw and Benjamini-Hochberg-adjusted p-values were reported for each gene.

#### 4.6.3. Pathway analysis

Log2 fold-changes or Wald’s statistics were retrieved from DESeq2 runs used as a ranking for Gene Set Enrichment Analysis (GSEA) 3.0 against MsigDB collections c2, c3, c4, c5 and c6 v.7.0,^[73,74]^ with flags -nperm 5000 -set_min 5 -set_max 2000 -plot_top_x 1000.

### 4.7. Immunofluorescence Imaging

During the duration of the experiment, live-cell imaging was performed using an EVOS fluorescence. After live-cell experiments were performed, tissues were fixed in 4% paraformaldehyde in PBS, permeabilized using 0.5% Triton X-100 in PBS (Sigma-Aldrich, X100), and blocked for an hour in antibody diluent (Thermofisher, 003218). Samples were incubated with primary antibody, MYH4 (1:200), in antibody diluent at 4° C overnight. Then, the samples were rinsed with PBS three times before being incubated with the secondary antibody, Alexa Fluor 488 goat anti-rabbit (1:1000), for an hour at room temperature. Finally, the samples were rinsed with PBS three times before and after staining the nuclear DNA with NucBlue and Phalloidin-iFluor 647(Abcam, ab176759) for 30 minutes at room temperature.

### 4.8. Neuron Morphological Data Analysis

The morphological data was produced by analyzing the immunofluorescence images using ImageJ and NeuronJ, a plug-in for neurite analysis. Afterwards, the data was visualized and statistically analyzed using GraphPad Prism.

### 4.9 Muscle Displacement Tracking and Force Calculations

Muscle contraction videos were processed using our previously developed open-source computational framework (https://github.com/HibaKob/Raman_Manuscript_2023).^[1]^ Absolute displacement of pixels within a region of interest across subsequent image frames were calculated and plotted. The force generated by contracting muscle (F_muscle_) was calculated using the shear modulus of the fibrin (G ∼ G’ = 0.1 kPa for low-frequency contraction, as determined via rheological analysis), the displacement of the muscle during contraction (*x*), and known geometric parameters of the 2.5D tissue:

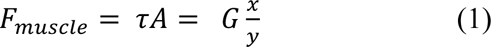

Where τ is the shear stress on the fibrin generated by the contractile muscle, *A* is the surface area of the fibrin (190 mm^2^), and *y* is the thickness of the fibrin (1 mm).

### 4.10. Gelatin Substrate Experiments

The C2C12 myoblasts were cultured to the procedure stated above, seeded onto the substrate in GM+ for 48 hours and then swapped onto DM++ for the remaining days. The gelatin substrate was fabricated in four different methods, varying between 0.5% w/v and 2% w/v of gelatin in PBS and varying between 5 and 30 minutes of incubation at 37° C before aspirating the gelatin solution. After fabricating the gelatin substrates on a glass-bottom well plate the C2C12s were seeded onto the substrate to begin the experiment. On day 7, the samples were fixed, permeabilized and blocked. We then stained and imaged the samples as mentioned in the procedure above.

## Acknowledgements

This work was supported by the U.S. DoD Army Research Office Early Career Program (awarded to R.R.), the NSF CAREER Program (awarded to R.R.), the PhRMA Foundation Research Starter Grant in Translational Medicine (awarded to R.R.), the MIT Center for Multicellular Engineered Living Systems seed grant (awarded to R.R.), and the NSF Graduate Research Fellowship Program (awarded to A.B. and B.R.). Our research was partly conducted within the Koch Institute’s Robert A. Swanson (1969) Biotechnology Center, supported by a core grant P30-CA14051 from the National Cancer Institute, and benefited from technical support from the Peterson (1957) Nanotechnology Materials Core Facility (RRID:SCR_018674).

