## Supplementary Information for "2.5D Actuating Substrates Enable Decoupling the Mechanical and Biochemical Effects of Muscle Exercise on Motor Neurons"

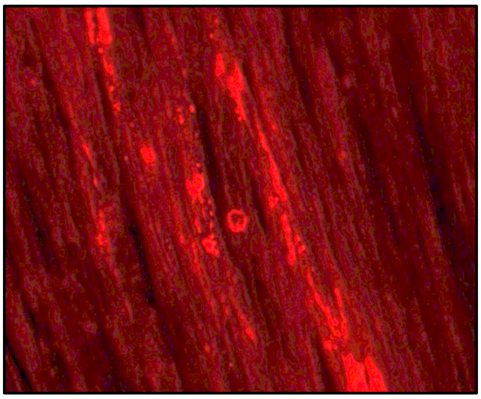

**Video S1.** Spontaneous twitch of 2.5D muscle tissue on fibrin substrates.

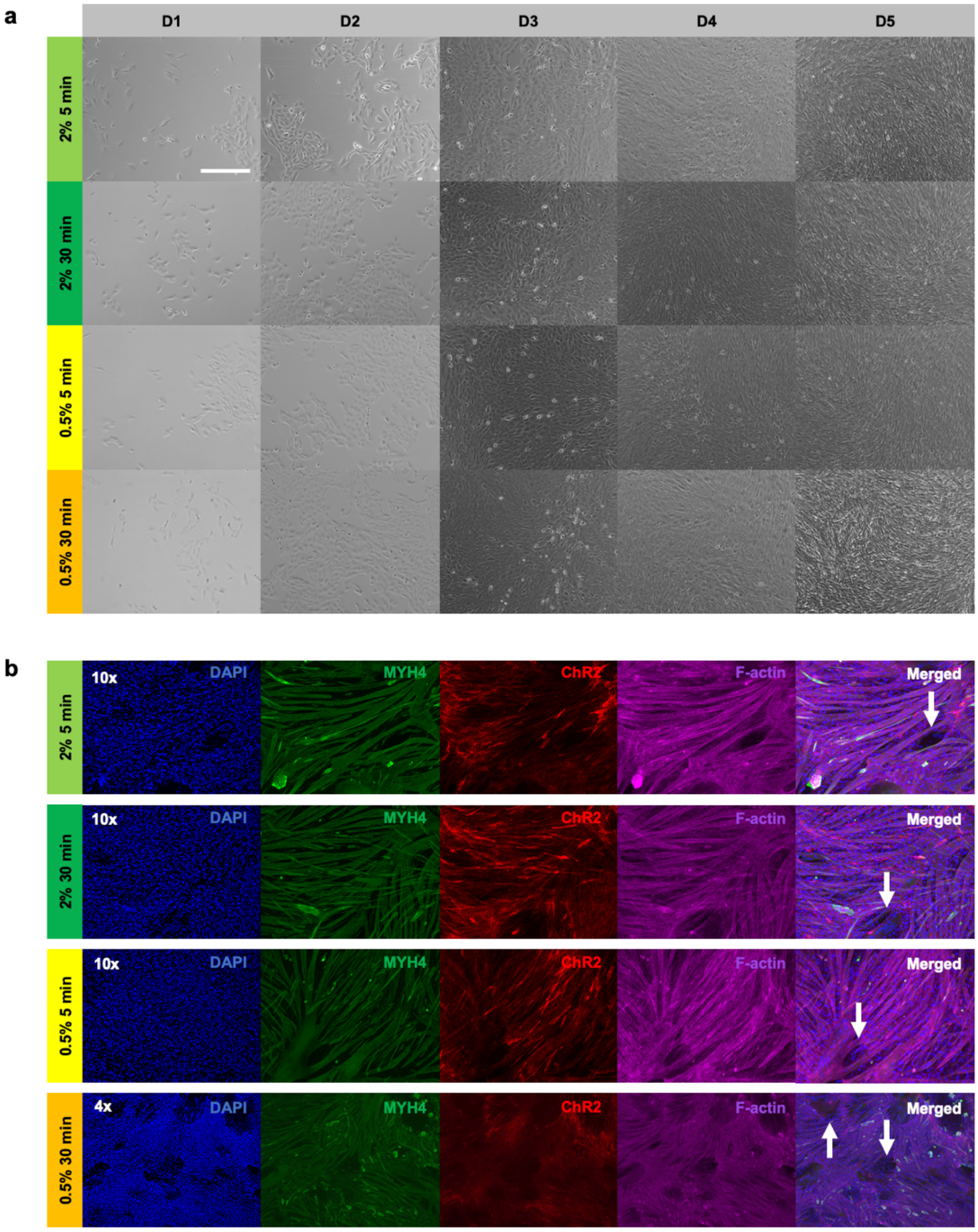


**Figure S1. Muscle monolayer cultured on different gelatin substrates. a)** Representative images of our optogenetic C2C12 myoblasts growing on different gelatin substrates. The gelatin substrates included (1$^{st}$ row) 2% w/v of gelatin incubated for 5 minutes, (2$^{nd}$ row) 2% w/v of gelatin incubated for 30 minutes, (3$^{rd}$ row) 0.5% w/v of gelatin incubated for 5 minutes, and (4$^{th}$ row) 0.5% w/v gelatin incubated for 30 minutes. Scale bar = 300 μm. **b)** Immunofluorescence images of the muscle monolayer on day 7 in the same order of substrates as above. The muscle monolayer is shown using DAPI (nuclei), MYH4 (myosin), ChR2 (optogenetic ion channel), F-actin (cytoskeleton), and merged channels. The muscle layer has delaminated in several locations on each of the substrates, as indicated by the white arrows showing a few black regions with no cells.

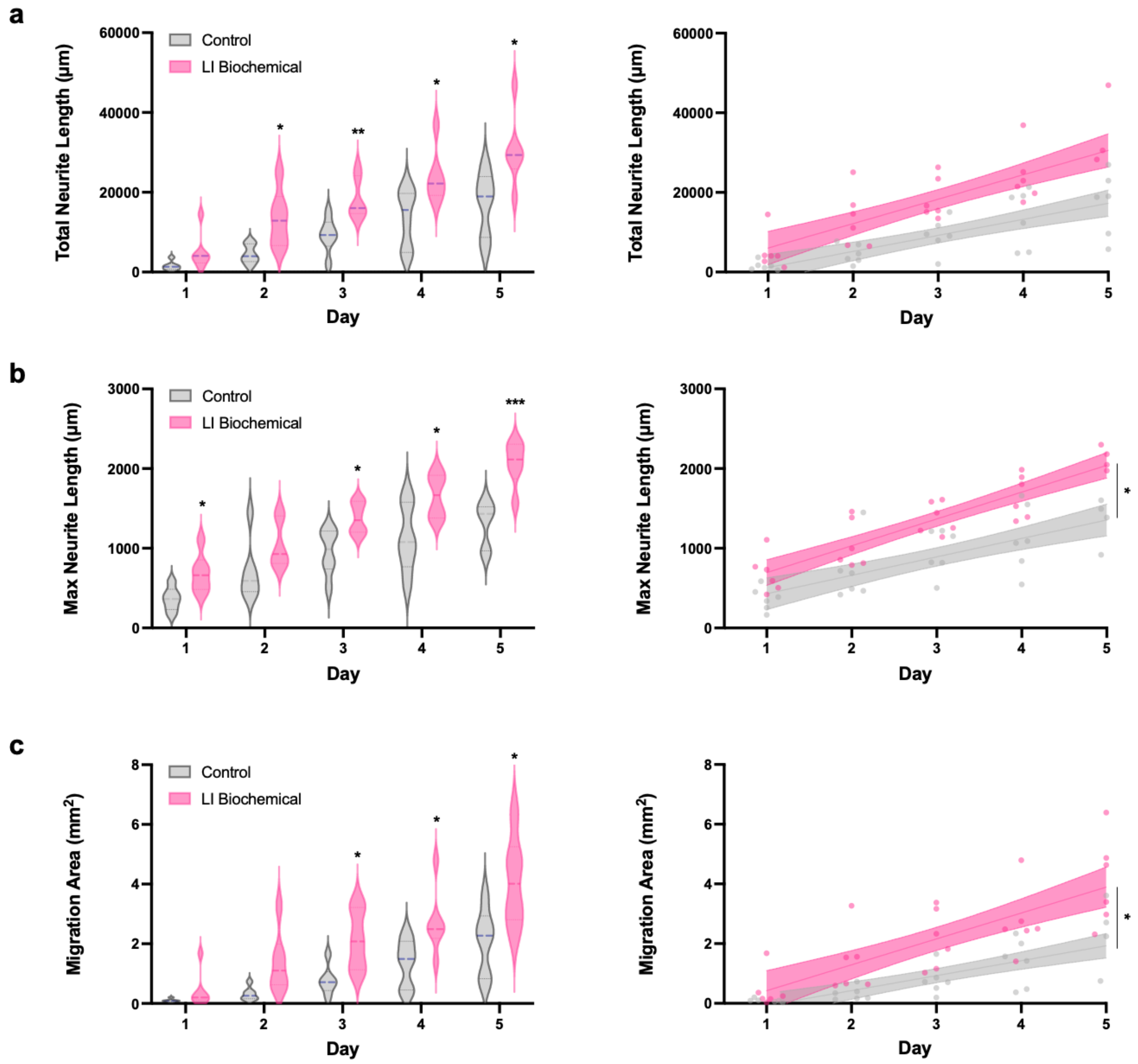


**Figure S2. Impact of low intensity biochemical stimulation on motor neuron growth.** Low intensity biochemical stimulation was performed by supplementing motor neuron spheroids with conditioned media from spontaneously twitching muscle. Low intensity biochemical stimulation was observed to significantly increase the **a)** total neurite length, **b)** max neurite length, and **c)** migration area in the later stages of the experiment. The rate of growth for max neurite length and migration area were also significantly impacted by biochemical stimulation. Unpaired t-test, n = 6 per group, $p<0.05$*, $p<0.01$**, $p<0.001$***.
